# Rift Valley fever virus minigenome system for investigating the role of L protein residues in viral transcription and replication

**DOI:** 10.1101/556738

**Authors:** Hanna Jérôme, Martin Rudolf, Michaela Lelke, Meike Pahlmann, Carola Busch, Sabrina Bockholt, Stephanie Wurr, Stephan Günther, Maria Rosenthal, Romy Kerber

## Abstract

Replicon systems are important molecular tools for investigating the function of virus proteins and regulatory elements involved in viral RNA synthesis. Various such systems were previously established for segmented negative strand viruses including the Rift Valley fever virus (RVFV). We have developed an ambisense minigenome system for RVFV with the specific aim to analyze the effects of L gene mutations on viral transcription versus replication. The S RNA segment with regulatory elements for ambisense gene expression served as backbone for the minigenome. Expression of the luciferase reporter gene allowed the overall activity of the RVFV replication complex to be assessed, while northern blot analysis enabled differentiation between synthesis of viral mRNA and replication intermediates. The functionality of the system was demonstrated by probing residues predictably involved in the active site of the cap-snatching endonuclease in the N-terminus of the L protein (D111, E125, and K143). Corresponding mutations led to a selective defect in the viral mRNA synthesis as described for other viruses of the *Bunyavirales* order. The analysis of further L gene mutants revealed an essential and specific role of a C-terminal region in the RVFV L protein (residues 1680–2068) in viral transcription. In summary, the established minigenome system is suitable for functional testing of the relevance of residues for viral transcription and replication and to validate hypotheses arising from structural or biochemical investigations of the RVFV replication complex. Application of the system to a small-scale mutagenesis screen disclosed a specific role of a C-terminal region in the RVFV L protein in mRNA synthesis.

## Introduction

Rift Valley fever virus (RVFV) is an important human and animal pathogen in Sub-Saharan Africa and on the Arabian Peninsula. RVFV infections cause death and abortion in ruminants and pseudoruminants and outbreaks may be associated with a high economic burden. In humans, the virus may cause febrile illness including hemorrhagic fever with fatal outcome [1, 2]. As medical countermeasures to prevent or treat the disease in humans are lacking, it is listed on the WHO blueprint for urgent research and development [3].

RVFV belongs to the family of *Phenuiviridae* within the *Bunyavirales* order and contains a trisegmented single stranded RNA genome with negative polarity. The small (S) RNA segment utilizes an ambisense coding strategy; the nucleocapsid protein (N) is encoded in antisense orientation and the non-structural protein (NSs) in sense orientation. Both genes are separated by an intergenic region (IGR). The middle and large (M and L) RNA segments contain genes for the glycoprotein precursor, the nonstructural protein NSm, and the large L protein (~250 kDa), respectively. N and L proteins together with the viral RNA constitute the viral replication complex, the structural unit for genome replication and transcription [4].

Several minigenome systems have been established for RVFV and were used to study the function of proteins and regulatory elements involved in RVFV genome replication and transcription [5–9]. However, these systems have not been optimized for comprehensive mutagenesis studies aiming at measuring simultaneously genome replication and transcription: (i) the measurement range (i.e. the ratio between positive and negative control) of the reporter gene assay is rather small [7, 9], although performance could be improved upon depletion of cellular protein kinase R [9], (ii) the read-out requires complex experimental procedures (i.e. virus-like particle production and transfer on indicator cells, chloramphenicol acetyltransferase [CAT] assay, and/or infection with Modified Vaccinia virus Ankara expressing T7 RNA polymerase [MVA-T7]) [5, 6, 8, 9], and/or (iii) the systems require transfection of plasmids hampering generation of larger numbers of L gene mutants [6–9].

In this article, we report the establishment of an ambisense minigenome system for RVFV based on the viral genomic S segment. It has been designed in analogy to the T7 RNA polymerase-driven minigenome system published for Lassa virus [10]. It provides a measurement range of two to three log units even after transfection of PCR products for expression of L protein, which is important for rapid and large-scale mutagenesis of the L gene. Luciferase reporter gene expression allows for assessment of the overall activity of the RVFV replication complex, while northern blot analysis of viral RNA facilitates easy discrimination between products of viral transcription and replication. A mutagenesis study for the L protein was conducted to demonstrate that the established RVFV system is suitable for studies with the aim of dissecting the molecular mechanisms of replication and transcription.

## Materials and methods

### Direct sequencing of RVFV

Overlapping fragments of RVFV-ZH501-BNI S, M, and L RNA, respectively, were reverse transcribed, amplified, and sequenced directly. In order to generate an S RNA based minigenome with an authentic promoter of strain ZH501-BNI, the conserved 5’ and 3’ termini of S RNA were sequenced as described previously [11]. In brief, purified virus RNA was treated with 5 units of tobacco acid pyrophosphatase (Epicentre) to generate 5’ monophosphorylated termini. Subsequently 5’ and 3’ termini were ligated using 10 units of T4 RNA ligase (New England Biolabs) at 37°C for 1 h. The resulting intramolecular ligation site was reverse transcribed and amplified, and the PCR product was sequenced. Sequences of primers used can be obtained upon request. Compared to the sequences of RVFV strain ZH501 deposited in GenBank (accession numbers DQ375406, DQ380200, DQ380149), the S, M and L RNA sequences for RVFV strain ZH501-BNI show the following nucleotide (amino acid) differences: S RNA, C616A (NSs protein: C194Stop); M RNA, A715T (envelope polyprotein: Q232L), T895C (envelope polyprotein: L292P), G3614A (envelope polyprotein: Stop1198Stop).

### Construction of plasmids for RVFV minigenome system

Vero E6 cells in 75-cm^2^ tissue culture flasks were inoculated with RVFV-ZH501-BNI. After 4 days, the supernatant was cleared by low-speed centrifugation and virus was pelleted by overnight ultracentrifugation. The pellet was resuspended in water, and virus RNA was purified by using the QIAamp viral RNA kit (Qiagen) according to the manufacturer’s instructions. Purified RNA was reverse transcribed and the resulting cDNA amplified using the Superscript III One-Step RT-PCR System with Platinum Taq (Invitrogen). Amplified RVFV genes for N and L proteins were cloned into expression vector pCITE-2a containing a T7 RNA polymerase promoter, an internal ribosomal entry site (IRES), and a favorable Kozak consensus sequence, resulting in pCITE-RVFV-N and pCITE-RVFV-L, respectively. The final sequences of N and L genes in pCITE 2a vector matched the consensus sequence of strain ZH501-BNI. The RVFV minigenome plasmid (pRVFV-MG) is based on the genomic RVFV S RNA integrated into the vector pX12ΔT [12]. The pRVFV-MG contains the T7 RNA polymerase promoter followed by a single G base, 5’ untranslated region (5’-UTR) including the conserved 5’ terminus, CAT gene, 3’ end of the NSs gene (55 nucleotides), the IGR, 3’ end of the N gene (49 nucleotides in reverse orientation), Renilla luciferase (Ren-Luc) gene in reverse orientation, 3’-UTR including the conserved 3’ terminus, hepatitis delta ribozyme (HDR), and T7 RNA polymerase transcription termination sequence (T7t) (Fig 1). The sequence of RVFV IGR and flanking nucleotides of the N and NSs genes were synthesized by GeneArt (Thermo Fisher) for stepwise assembly of the complete minigenome in the pX12ΔT vector, resulting in pRVFV-MG. Correctness of all sequences was ascertained by sequencing. Sequences of primers used for cloning are available upon request.

**Fig 1.**
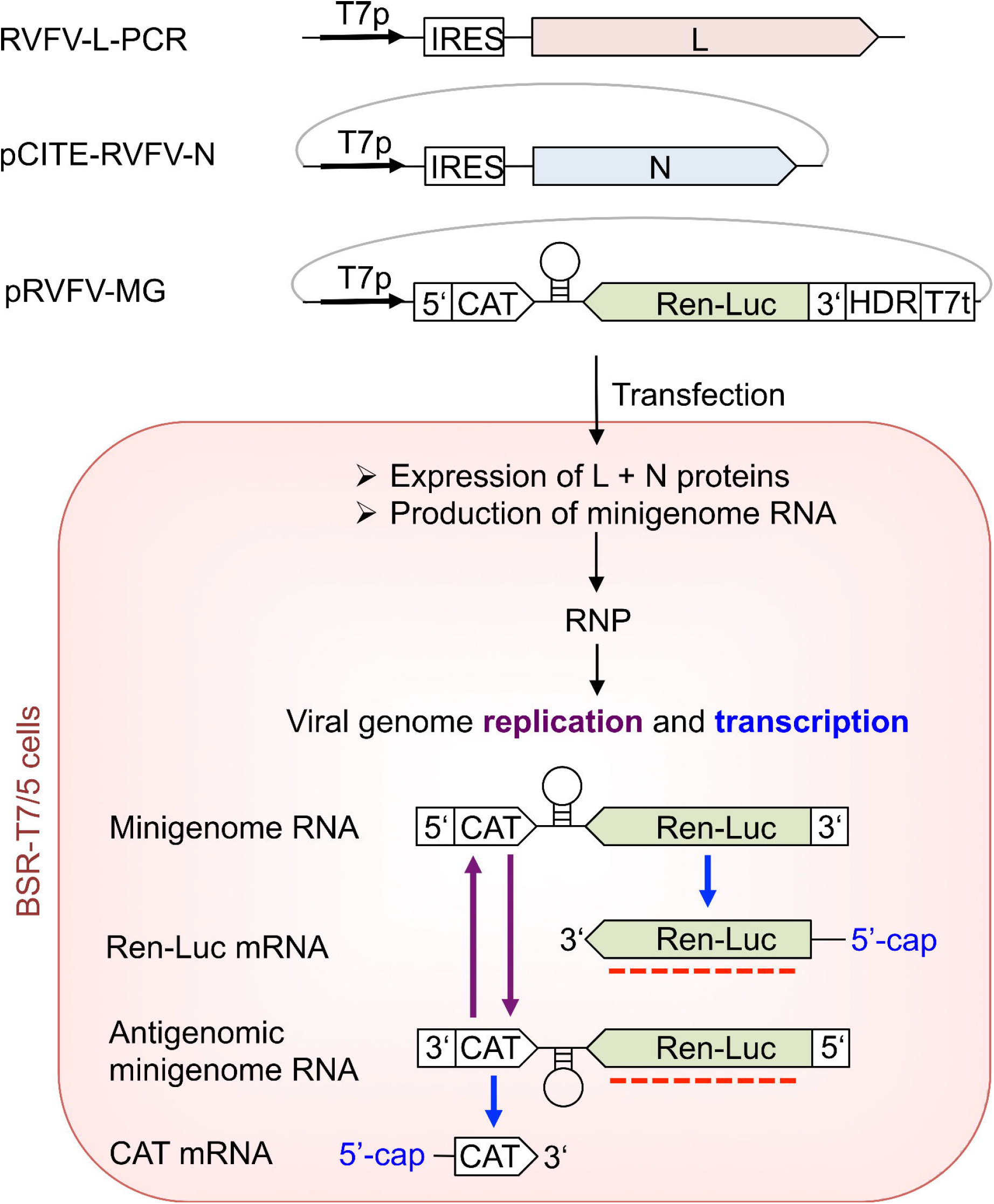
The RVFV ambisense minigenome system. The upper part of the figure schematically displays the minigenome system constructs as used for transfection into BSR-T7/5 cells. The system comprises the RVFV minigenome plasmid pRVFV-MG based on RVFV S RNA, the plasmid for expression of the N protein (pCITE-RVFV-N), and the PCR products for expression of the L protein (RVFV-L-PCR). Plasmid pCITE-FF expressing firefly luciferase serves as a transfection control (not depicted). Functional elements are abbreviated as follows: T7p, T7 RNA polymerase promoter; IRES, internal ribosmal entry site; UTR, untranslated region with conserved termini of the open reading frames of the N and the NSs genes; CAT, chloramphenicol acetyltransferase gene; IGR, intergenic region, Ren-Luc, Renilla luciferase gene; HDR, hepatitis delta ribozyme; T7t, T7 RNA polymerase transcription termination sequence. These constructs are transfected into BSR-T7/5 cells and lead to production of L and N proteins as well as minigenome RNA, which are the minimal components for viral replication and transcription and form viral ribonucleoparticles (RNP). By the processes of viral genome replication antigenomic minigenome RNA and minigenome RNA are produced. Ren-Luc and CAT mRNAs are transcribed from the minigenome RNA and antigenomic minigenome RNA, respectively. The mRNAs contain 5’-cap structures obtained by the cap-snatching mechanism. A red dotted line indicates the targets of the riboprobe used for the northern blot analysis.

### Minigenome assay

Mutant L genes were generated by mutagenic PCR using pCITE-RVFV-L as a template. The PCR products containing the functional cassette for expression of mutant L protein (T7 RNA polymerase promoter, IRES, and L gene) were purified, quantified spectrophotometrically, and used for transfection without prior cloning as described previously [13]. The presence of the artificial mutation was ascertained by sequencing. BSR-T7/5 cells stably expressing T7 RNA polymerase [12] were transfected per well of a 24-well plate with 250 ng of L gene PCR product, 750 ng of pRVFV-MG expressing Renilla luciferase (Ren-Luc), 500 ng of pCITE-RVFV-N expressing N protein, and 10 ng of pCITE-FF-Luc expressing firefly luciferase as an internal transfection control. All transfections were performed by use of Lipofectamine 2000 (Thermo Fisher) according to the manufacturer’s instructions. At 24 h after transfection, cells were either used to purify total RNA using the RNeasy Mini Kit (Qiagen) for northern blotting, or lysed in 100 μl of passive lysis buffer (Promega) per well, and analysed for firefly luciferase and Ren-Luc activity using the Dual-Luciferase Reporter Assay System (Promega). To compensate for differences in transfection efficiency or cell density, Ren-Luc levels were corrected with the firefly luciferase levels (resulting in standardized relative light units [sRLU]). RNA (1–2 μg) was separated in a 1.5%–agarose-formaldehyde gel and transferred onto a Roti-Nylon Plus membrane (Roth) for northern blot analysis. After pre-hybridization, blots were hybridized with a ^32^P-labeled riboprobe targeting the Ren-Luc gene (Fig 1). RNA bands were visualized by autoradiography using a Typhoon scanner (GE Healthcare).

### Expression of L protein

To verify expression of L protein mutants, BSR-T7/5 cells in a well of a 24-well plate were transfected with 500 ng of PCR product encoding L protein mutants tagged at the C-terminus with a 3xFLAG sequence. Cells were additionally inoculated with MVA-T7 [14] prior to the transfection to enhance L protein expression levels. At 24 h after transfection, cytoplasmic lysate was separated in a 3–8% Tris-acetate polyacrylamide gel, transferred to a nitrocellulose membrane (Whatman), and detected by immunoblotting using peroxidase-conjugated anti-FLAG M2 antibody (1:10,000) (A8592; Sigma-Aldrich). For visualization of the L protein bands by chemiluminescence, the SuperSignal West Femto substrate (Pierce) and a FUSION SL image acquisition system (Vilber Lourmat) were used.

## Results and discussion

Essential components for an ambisense minireplicon system for RVFV, namely L gene, N gene, and minigenome, were integrated into appropriate vectors for T7 RNA polymerase-driven expression in BSR-T7/5 cells (Fig 1). The S RNA segment containing regulatory elements for ambisense gene expression was used as backbone for the minigenome. The latter contained CAT and Ren-Luc genes in sense and antisense orientation, respectively, to measure transcriptional activity of the system, although only Ren-Luc expression, which is solely dependent on the activity of the RVFV polymerase, was measured in this study. Expression of firefly luciferase from cotransfected plasmid served as internal control for transfection efficacy. To demonstrate functionality of the system, experiments were conducted with wild-type L protein. An L protein mutant containing a mutation in the catalytic site of the RNA-dependent RNA polymerase (D1133N within the SDD motif) served as a negative control. Wild-type L protein mediated high levels of Ren-Luc expression (up to 1,500,000 light units), while the inactive mutant showed 100–1,000 fold less Ren-Luc expression. RNA products of genome replication (antigenome) and transcription (Ren-Luc mRNA) generated by the wild-type L protein were clearly visible as distinct signals in northern blot (Fig 2 and S1 Table), similar to the ambisense Lassa virus minigenome system [10, 15, 16]. Transcription signals for the D1133N mutant were absent but an unspecific background signal at the antigenome position was sometimes observed on the northern blot, which was taken into account in the quantification of antigenome signal intensity. The precise nature of the unspecific material is not known, although it likely stems from spurious activity of cellular enzymes, which use either the transfected pRVFV-MG plasmid or the RNA expressed from this plasmid by T7 RNA polymerase as a template for synthesis of an “antigenome-like” RNA species. The activity of such cellular enzymes may also explain background expression of Ren-Luc in the absence of a functional RVFV L protein.

**Fig 2.**
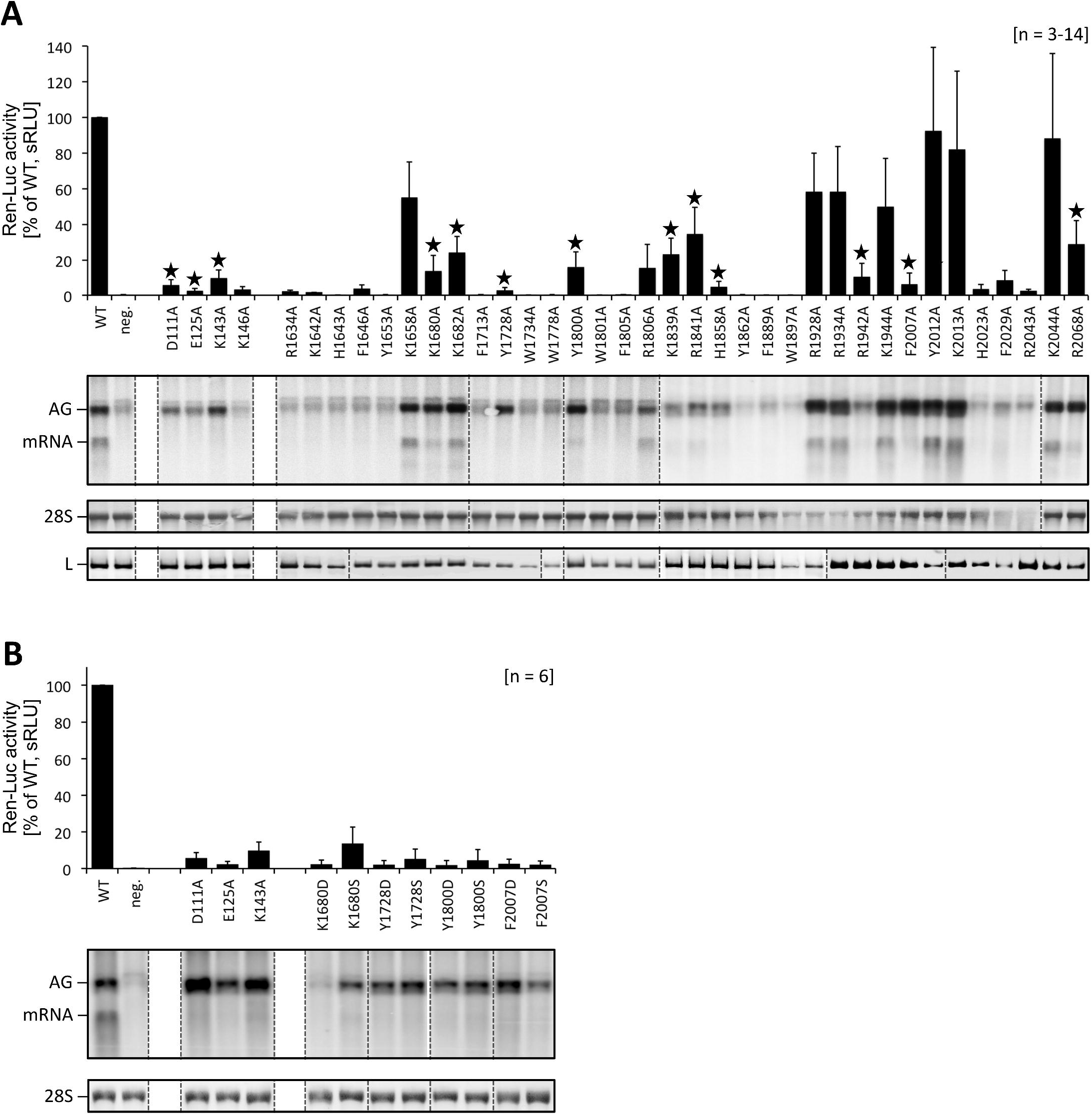
Determination of transcription and replication activity of L protein mutants using the RVFV ambisense minigenome system. **(A)** The activity of L protein mutants in viral replication and transcription was measured via Ren-Luc reporter gene expression. The Ren-Luc activity is shown in the bar graph (mean and standard deviation of standardized relative light units [sRLU] as a percentage of the wild-type (WT) in 3 to 14 independent transfection experiments). Signals for antigenomic RNA (position AG, representing viral replication) and Ren-Luc mRNA (position mRNA, representing viral transcription) were detected by northern blotting using a radiolabeled riboprobe hybridizing to the Ren-Luc gene. A defective L protein with a mutation in the catalytic site of the RNA-dependent RNA polymerase (D1133N) served as a negative control (neg.). Northern blots were performed two to three times per mutant and signals on northern blots were quantified using ImageJ2 software [20]. The quantitative data are presented in S1 Table. The methylene blue-stained 28S rRNA (28S) served as a marker for gel loading and RNA transfer. Additionally, immunoblot analysis of 3xFLAG-tagged L protein mutants is shown (L). Mutants with an mRNA defective phenotype are marked with an asterisk. For experimental details see methods section. Dotted lines indicate removal of irrelevant lanes for presentation purposes. Original blots are included in the supporting information (S1 File). **(B)** Further L protein mutants were tested in the RVFV ambisense minigenome system essentially as described in (A). The bar graph represents mean and standard deviation of 6 independent measurements for the Ren-Luc activity. Northern blot analysis was performed twice; the figure depicts one representative experiment. The quantitative data are presented in S1 Table. Dotted lines indicate removal of irrelevant lanes for presentation purposes. Original blots are included in the supporting information (S1 File).

In order to validate the RVFV minigenome system, we tested L protein mutants with exchanges of residues presumably involved in the endonuclease active site. The endonuclease domain has been located in the N-terminal ~250 residues of RVFV L protein [9] and is predictably required for viral transcription, as demonstrated for the corresponding domain of Lassa virus L protein [15, 17]. Putative catalytic residues D111, E125 and K143 were selected based on amino acid alignments of RVFV with La Crosse virus and hantavirus endonuclease domains as well as prior functional and structural information on the catalytic site [9, 17–19]. Mutation of these residues to alanine resulted in a strong decrease in Ren-Luc activity (Fig 2 and S1 Table). On the northern blot, replication products were detected, whereas Ren-Luc mRNA was absent consistent with the low level of Ren-Luc activity. In summary, alanine substitution of RVFV L protein residues predictably involved in endonuclease activity led to a selective defect in viral transcription. The transcription-defective phenotype could be demonstrated using the ambisense minigenome system.

Besides the endonuclease, a cap-binding function is important for viral transcription. It has been hypothesized that the C terminus of bunyavirus L protein is involved in this function [21, 22]. Furthermore, a role for the C terminus of Lassa virus L protein in viral transcription was proposed based on a mutagenesis study using the Lassa virus minigenome system [16]. A typical structural motif of the cap-binding site comprises two aromatic amino acid side chains forming a sandwich with the guanine moiety of a cap structure. Additionally, the triphosphate moiety of the cap structure is often interacting with positively charged amino acids. However, a cap-binding site does not feature a specific sequence motif; therefore it is not possible to predict residues potentially involved in cap-binding just based on sequence.

We used the RVFV minigenome system to investigate whether the C-terminal region of RVFV L protein might play a role in viral cap-snatching. Based on an alignment of phlebovirus L protein sequences, 34 partially or completely conserved aromatic and positively charged amino acids were selected for an alanine mutation screen (S1 Fig). Ten of these residues were found to be important for transcription but not replication of the viral genome (Fig 2 and S1 Table). Four residues were aromatic or heteroaromatic (Y1728, Y1800, H1858 and F2007) and six were positively charged (K1680, K1682, K1839, R1841, R1942 and R2068). A subset of these residues was additionally modified to serine and aspartic acid. All these modifications resulted in a selective defect in mRNA synthesis, which confirmed the involvement of the C-terminal region between positions 1680 and 2068 of the RVFV L protein in viral transcription (Fig 2 and S1 Table).

We developed an ambisense minigenome system for RVFV, which is suitable for screening of L gene mutations for their impact on viral transcription and replication. As a proof of principle, the system was used to probe amino acid residues potentially involved in the endonuclease active site (D111, E125, and K143). Corresponding mutations resulted in a selective defect in viral transcription, as has been reported for other viruses of the *Bunyavirales* order [15, 23]. Additionally, the system facilitated identifying residues in the C-terminal region of RVFV L protein (residues 1680–2068) being important for viral transcription but not replication. However, these data is no proof for the existence of a cap-binding site. They merely demonstrate a specific role of the identified amino acids during viral transcription. Further conclusions require biochemical and structural data.

In summary, the established RVFV ambisense minigenome system (i) is suitable to screen L protein mutants without cloning, (ii) yields sufficient signal strength without depletion of the cellular protein kinase R [9], and (iii) allows for technically simple discrimination between viral transcription and replication. Therefore this system is well suited to validate hypotheses arising from structural or biochemical investigation of the RVFV replication complex.

## Author statements

### Author contributions

Conceptualization: SG; investigation: HJ, MRu, ML, MP, CB, SB, SW; supervision: SG, RK; data analysis: MRo, RK; visualization: MRo, RK; writing – original draft preparation: MRo, RK; writing –review & editing: SG, MRo, RK

## Acknowledgement

We thank Martin Meyer and Beate Becker-Ziaja for excellent technical assistance. We thank Sophia Reindl for fruitful discussions and also acknowledge support by Stephanie Jansen and Stefanie Becker.

## Supporting data

**S1 Table. Functional analysis of L protein mutants in the RVFV ambisense minigenome system.**

^1^ For each mutant, 3 to 14 independent transfection experiments were performed. Ren-Luc values represent mean with standard deviation (n = 3-14). Northern blots were performed at least twice for every mutant. A selective defect in mRNA synthesis was defined as reduction in Ren-Luc level (≤1–35%) despite wild-type like antigenome synthesis (37–280%) and reduction of the mRNA-to-antigenome ratio (≤1–35%). Mutants with a selective defect in mRNA synthesis are shown in boldface on grey background.

^2^ Standardized relative light unit (sRLU) value (wild-type L protein = 100%).

^3^ sRLU value of mutant divided by sRLU value of the negative control mutant (D1133N) containing a mutation in the catalytic site of the RNA-dependent RNA polymerase.

^4^ Antigenome signals in northern blots were quantified via intensity profiles using ImageJ2 software (wild-type L protein = 100%). Background signals at the position of the antigenome of the respective northern blot were either subtracted from all other replication signals or data were evaluated without background subtraction (numbers in parentheses).

^5^ RNA signals on northern blots were quantified using ImageJ2 software and the mRNA-to-antigenome signal ratio was calculated. The wild-type ratio was set at 100% for each experiment (i.e., the signal ratio of a mutant was normalized with the wild-type ratio) to render independent experiments comparable. Background signals at the position of the antigenome were subtracted from all other replication signals. Data without background correction are also shown (numbers in parentheses).

**S1 Figure. Alignment of C terminal sequences of phlebovirus L proteins.** The figure presents a secondary structure-guided alignment of the C-terminal sequences of L proteins from 22 phleboviruses (Uniprot accession numbers are given). The alignment was created by manually combining results of ClustalOmega and PRALINE programs [24–27] and data are presented by ESPript (http://espript.ibcp.fr) [28]. The corresponding secondary structure predictions were calculated by Jpred4 [29] and are depicted below the sequences (β-sheets as arrows, α-helices as barrels, loops as lines). All positions refer to RVFV strain ZH-501 full-length L protein. Impact of residue-to-alanine substitution on RVFV L protein activity is indicated by triangles above the sequences (white, mild or no effect; grey, general defect in L protein activity; red, selective defect in viral transcription).

**S1 File. Original northern and western blots.** Lanes of northern blots that have been used for quantification and statistical analysis (included in S1 Table) are labeled with the respective L protein mutation, “WT” for L protein wild-type, or “neg.” for negative control mutant D1133N. Lanes of northern blots and methylene blue stained northern blot membranes presented in Fig 2 are framed by dotted rectangles. Western blot bands used in Fig 2 are also framed by dotted rectangles and labeled.

