## Supporting data for "Rift Valley fever virus minigenome system for investigating the role of L protein residues in viral transcription and replication"

**S1 Table. Functional analysis of L protein mutants in the RVFV ambisense minigenome system.[1]**

| Mutant | Renilla luciferase activity (sRLU) | | | | RNA expression level (Northern blot signal) | |
|---|---|---|---|---|---|---|
| | % of wild-type[2] | | | Signal-to-noise ratio[3] | Antigenome level % of wild-type[4] | mRNA-to-antigenome ratio, %, relative to wild-type[5] |
| D1133N | ≤ 1 | +/– | 1.4 | 1.0 | 3.2 (14.7) | ≤ 1 (≤ 1) |
| **D111A** | **5.6** | **+/–** | **3.2** | **9.9** | **60.5 (92.4)** | **≤ 1 (≤ 1)** |
| **E125A** | **2.3** | **+/–** | **1.7** | **4.0** | **37.4 (75.7)** | **≤ 1 (≤ 1)** |
| **K143A** | **9.7** | **+/–** | **4.7** | **17.4** | **111.5 (116.4)** | **≤ 1 (≤ 1)** |
| K146A | 3.1 | +/– | 1.7 | 5.5 | 1.3 (49.6) | 57.8 (≤ 1) |
| R1634A | 2.2 | +/– | ≤ 1 | 3.9 | 10.4 (41.3) | ≤ 1 (≤ 1) |
| K1642A | 1.6 | +/– | ≤ 1 | 2.8 | 6.6 (38.4) | 1.72 (≤ 1) |
| H1643A | ≤ 1 | +/– | ≤ 1 | ≤ 1 | 13.4 (43.6) | ≤ 1 (≤ 1) |
| F1646A | 3.6 | +/– | 2.3 | 6.3 | 22.3 (50.4) | ≤ 1 (≤ 1) |
| Y1653A | ≤ 1 | +/– | ≤ 1 | ≤ 1 | 16.9 (46.2) | ≤ 1 (≤ 1) |
| K1658A | 54.9 | +/– | 20.0 | 97.8 | 222.9 (203.3) | 45.3 (47.1) |
| **K1680A** | **13.8** | **+/–** | **8.9** | **24.5** | **141.1 (138.7)** | **12.9 (12.8)** |
| **K1682A** | **24.0** | **+/–** | **9.3** | **42.7** | **248.9 (226.1)** | **17.4 (18.9)** |
| F1713A | ≤ 1 | +/– | ≤ 1 | ≤ 1 | 10.0 (48.2) | 76.2 (≤ 1) |
| **Y1728A** | **2.5** | **+/–** | **1.8** | **4.5** | **130.7 (129.5)** | **≤ 1 (≤ 1)** |
| W1734A | ≤ 1 | +/– | ≤ 1 | ≤ 1 | 23.1 (57.3) | ≤ 1 (≤ 1) |
| W1778A | ≤ 1 | +/– | ≤ 1 | ≤ 1 | 35.7 (66.2) | ≤ 1 (≤ 1) |
| **Y1800A** | **16.1** | **+/–** | **8.4** | **28.6** | **160.3 (154.4)** | **11.7 (11.7)** |
| W1801A | ≤ 1 | +/– | ≤ 1 | ≤ 1 | 27.4 (85.3) | ≤ 1 (≤ 1) |
| F1805A | ≤ 1 | +/– | ≤ 1 | ≤ 1 | 31.3 (90.4) | ≤ 1 (≤ 1) |
| R1806A | 15.4 | +/– | 13.5 | 27.4 | 65.1 (134.3) | 78.5 (82.4) |
| **K1839A** | **23.1** | **+/–** | **9.2** | **41.1** | **76.1 (80.3)** | **21.3 (19.1)** |
| **R1841A** | **34.5** | **+/–** | **15.0** | **61.4** | **59.4 (66.5)** | **21.4 (18.0)** |
| **H1858A** | **4.5** | **+/–** | **3.3** | **8.0** | **63.8 (70.1)** | **4.6 (4.1)** |
| Y1862A | ≤ 1 | +/– | ≤ 1 | ≤ 1 | ≤ 1 (24.2) | >100 (≤ 1) |
| F1889A | ≤ 1 | +/– | 0.07 | 0.29 | 10.2 (32.0) | 0.23 (≤ 1) |
| W1897A | ≤ 1 | +/– | 0.06 | 0.26 | 2.5 (26.1) | 0.95 (≤ 1) |
| R1928A | 58.2 | +/– | 21.9 | 103.6 | 228.1 (197.1) | 30.6 (26.9) |
| R1934A | 58.2 | +/– | 25.4 | 103.6 | 192.5 (170.1) | 39.1 (33.5) |
| **R1942A** | **10.5** | **+/–** | **7.7** | **18.7** | **69.0 (76.5)** | **18.4 (12.6)** |
| K1944A | 49.9 | +/– | 27.2 | 88.8 | 246.4 (210.9) | 26.2 (23.2) |
| **F2007A** | **6.1** | **+/–** | **6.8** | **10.9** | **191.5 (175.5)** | **4.3 (4.3)** |
| Y2012A | 92.4 | +/– | 47.0 | 164.5 | 275.8 (233.2) | 46.7 (41.9) |

| | | | | | | |
|---|---|---|---|---|---|---|
| K2013A | 82.0 | +/– | 44.0 | 146.1 | 311.2 (260.0) | 34.7 (31.5) |
| H2023A | 3.4 | +/– | 2.6 | 6.1 | 14.0 (29.1) | 12.4 (5.8) |
| F2029A | 8.2 | +/– | 6.0 | 14.6 | 26.3 (39.2) | 17.6 (10.6) |
| R2043A | 2.4 | +/– | ≤ 1 | 4.3 | 16.9 (31.5) | 4.3 (2.3) |
| K2044A | 88.2 | +/– | 47.7 | 157.0 | 275.1 (229.0) | 41.6 (36.8) |
| **R2068A** | **28.8** | **+/–** | **13.4** | **51.3** | **111.1 (109.2)** | **20.8 (18.8)** |
| **D111A** | **5.6** | **+/–** | **3.2** | **31.1** | **235.7 (159.3)** | **4.2 (5.1)** |
| **E125A** | **2.3** | **+/–** | **1.7** | **12.6** | **178.2 (120.5)** | **4.5 (5.0)** |
| **K143A** | **9.7** | **+/–** | **4.7** | **54.3** | **263.7 (178.3)** | **8.6 (11.0)** |
| **K1680D** | **2.3** | **+/–** | **2.5** | **12.5** | **48.4 (82.4)** | **>100 (16.0)** |
| K1680S | 13.5 | **+/–** | 9.3 | 75.1 | 124.5 (32.7) | 39.9 (33.2) |
| **Y1728D** | **1.9** | **+/–** | **2.4** | **10.7** | **169.6 (84.2)** | **5.6 (6.1)** |
| **Y1728S** | **5.2** | **+/–** | **5.6** | **28.9** | **201.6 (104.6)** | **7.4 (8.6)** |
| **Y1800D** | **1.9** | **+/–** | **2.5** | **10.4** | **179.9 (114.6)** | **9.7 (11.0)** |
| **Y1800S** | **4.5** | **+/–** | **6.0** | **25.0** | **159.2 (136.2)** | **14.9 (15.3)** |
| **F2007D** | **2.5** | **+/–** | **2.6** | **13.7** | **260.6 (81.9)** | **9.0 (11.3)** |
| **F2007S** | **1.9** | **+/–** | **2.3** | **10.6** | **168.9 (121.6)** | **11.7 (12.8)** |

[1] For each mutant, 3 to 14 independent transfection experiments were performed. Ren-Luc values represent mean with standard deviation (n = 3-14). Northern blots were performed at least twice for every mutant. A selective defect in mRNA synthesis was defined as reduction in Ren-Luc level (≤1–35%) despite wild-type like antigenome synthesis (37–280%) and reduction of the mRNA-to-antigenome ratio (≤1–35%). Mutants with a selective defect in mRNA synthesis are shown in boldface on grey background.

[2] Standardized relative light unit (sRLU) value (wild-type L protein = 100%).

[3] sRLU value of mutant divided by sRLU value of the negative control mutant (D1133N) containing a mutation in the catalytic site of the RNA-dependent RNA polymerase.

[4] Antigenome signals in northern blots were quantified via intensity profiles using ImageJ2 software (wild-type L protein = 100%). Background signals at the position of the antigenome of the respective northern blot were either subtracted from all other replication signals or data were evaluated without background subtraction (numbers in parentheses).

[5] RNA signals on northern blots were quantified using ImageJ2 software and the mRNA-to-antigenome signal ratio was calculated. The wild-type ratio was set at 100% for each experiment (i.e., the signal ratio of a mutant was normalized with the wild-type ratio) to render independent experiments comparable. Background signals at the position of the antigenome were subtracted from all other replication signals. Data without background correction are also shown (numbers in parentheses).

### S1 Figure

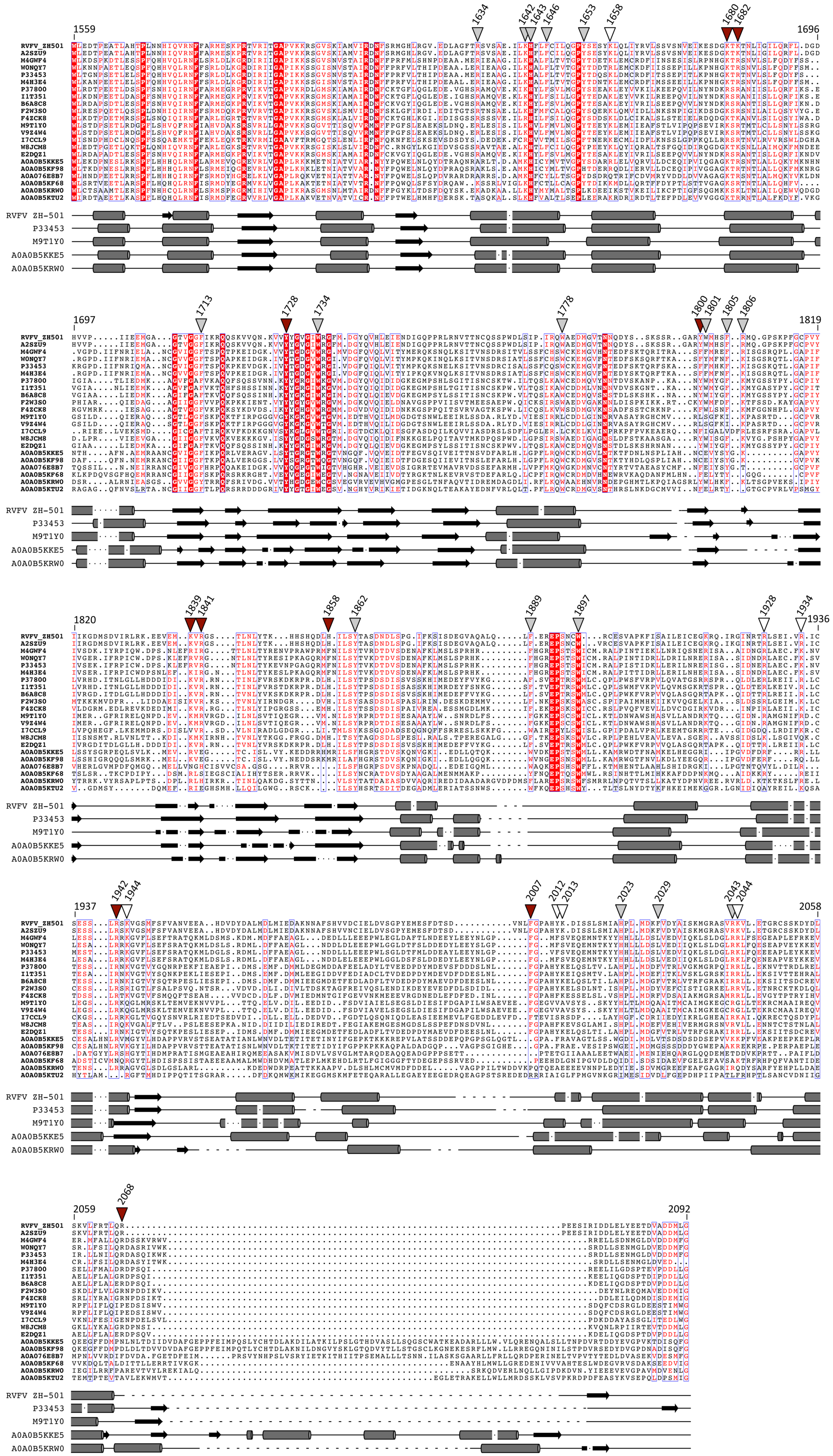
1559
1634
1642
1643
1646
1653
1658
1680
1682
1696
1697
1713
1728
1734
1778
1800
1801
1805
1806
1819
1820
1839
1841
1858
1862
1889
1897
1928
1934
1936
1937
1942
1944
2007
2012
2013
2023
2029
2043
2044
2058
2059
2068
2092
RVFV_ZH501
A2SZU9
M4GWF4
W0NQY7
P33453
M4H3E4
P37800
I1T351
B6A8C8
F2W3S0
F4ZCK8
M9T1Y0
V9Z4W4
I7CCL9
W8JCM8
E2DQZ1
A0A0B5KKE5
A0A0B5KF98
A0A076E8B7
A0A0B5KF68
A0A0B5KRW0
A0A0B5KTU2
RVFV ZH-501
P33453
M9T1Y0
A0A0B5KKE5
A0A0B5KRW0

S1 File

#### Northern Blot 1

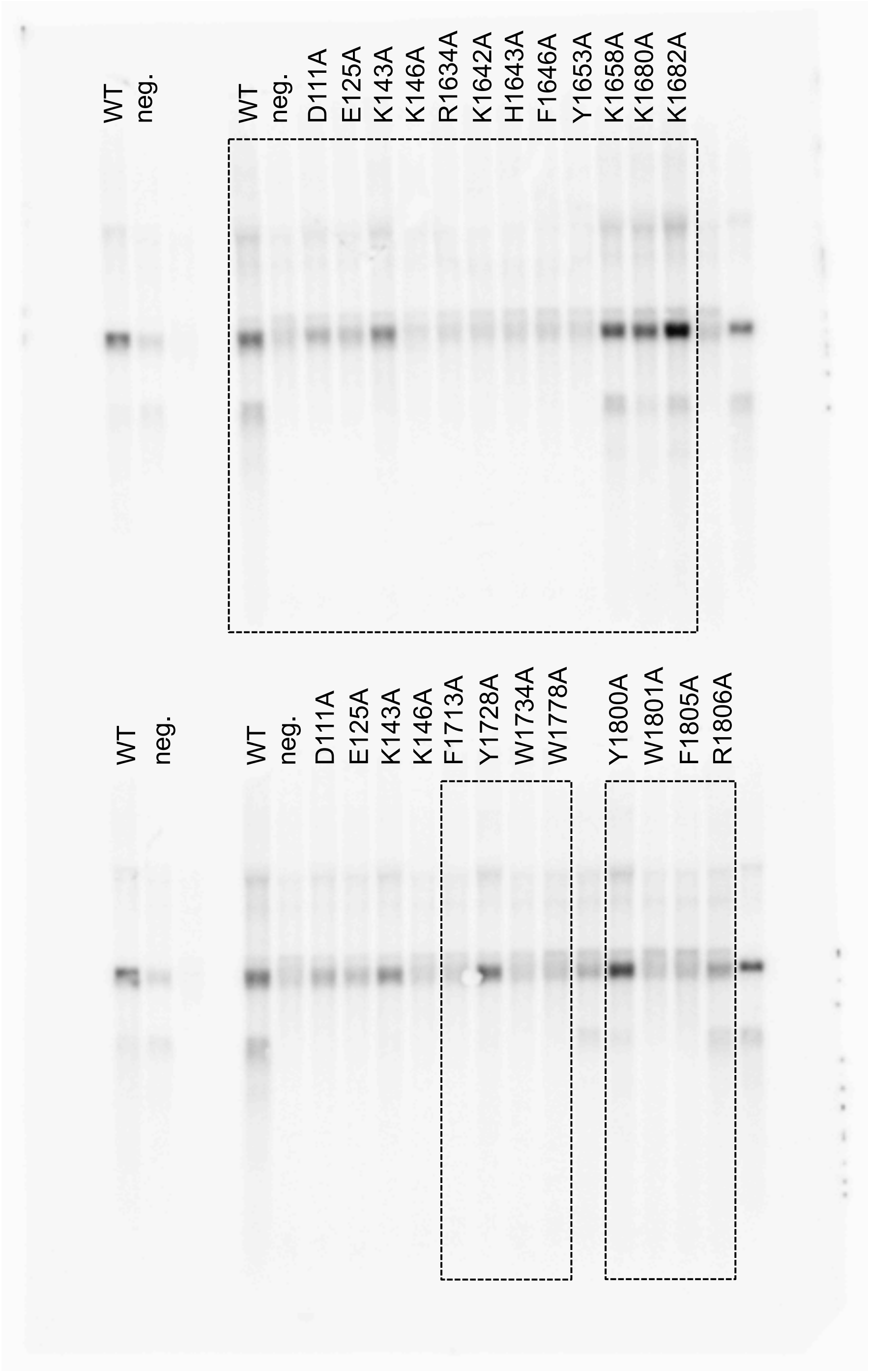

### Methylene blue staining of membrane 1

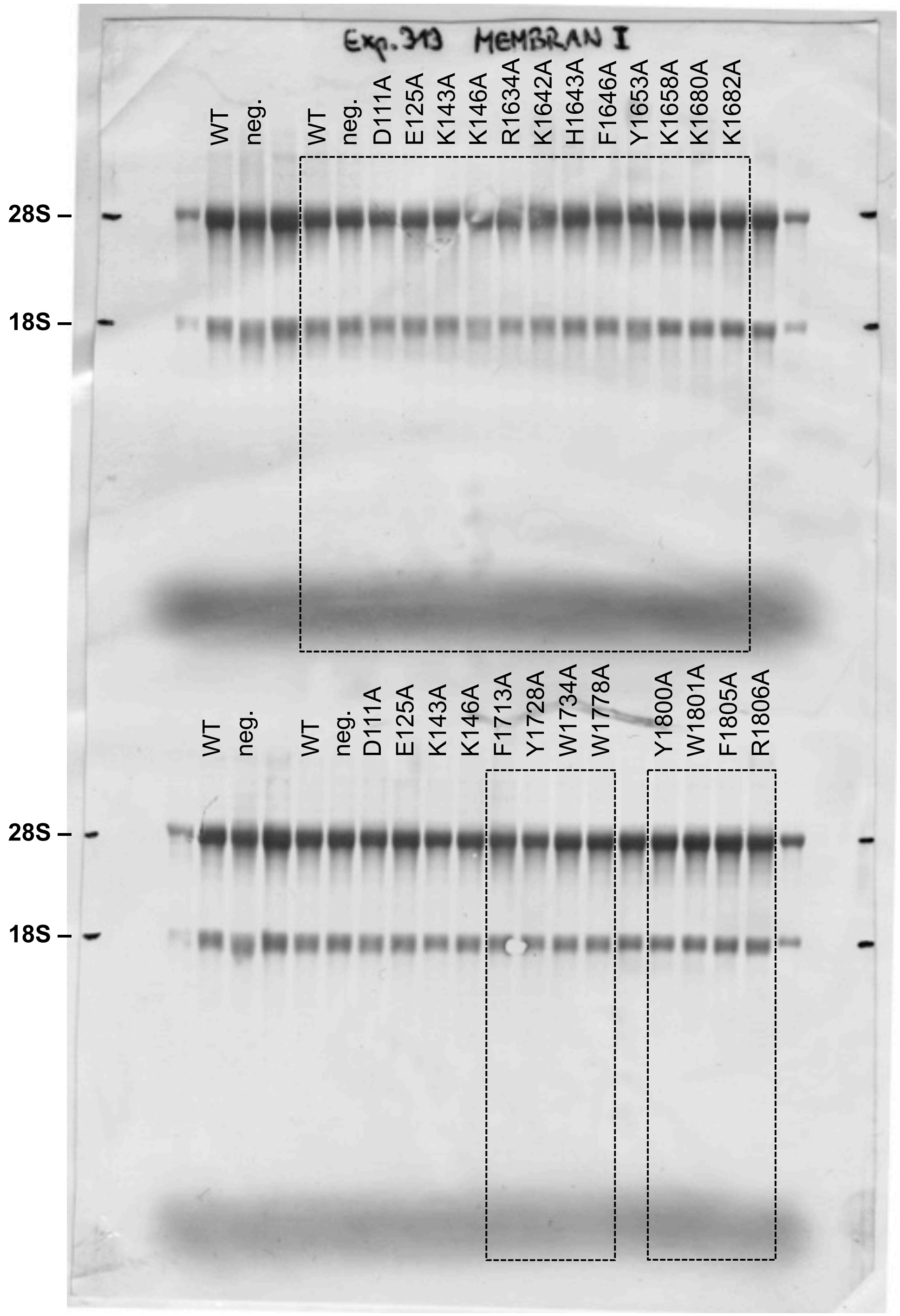

### Northern Blot 2

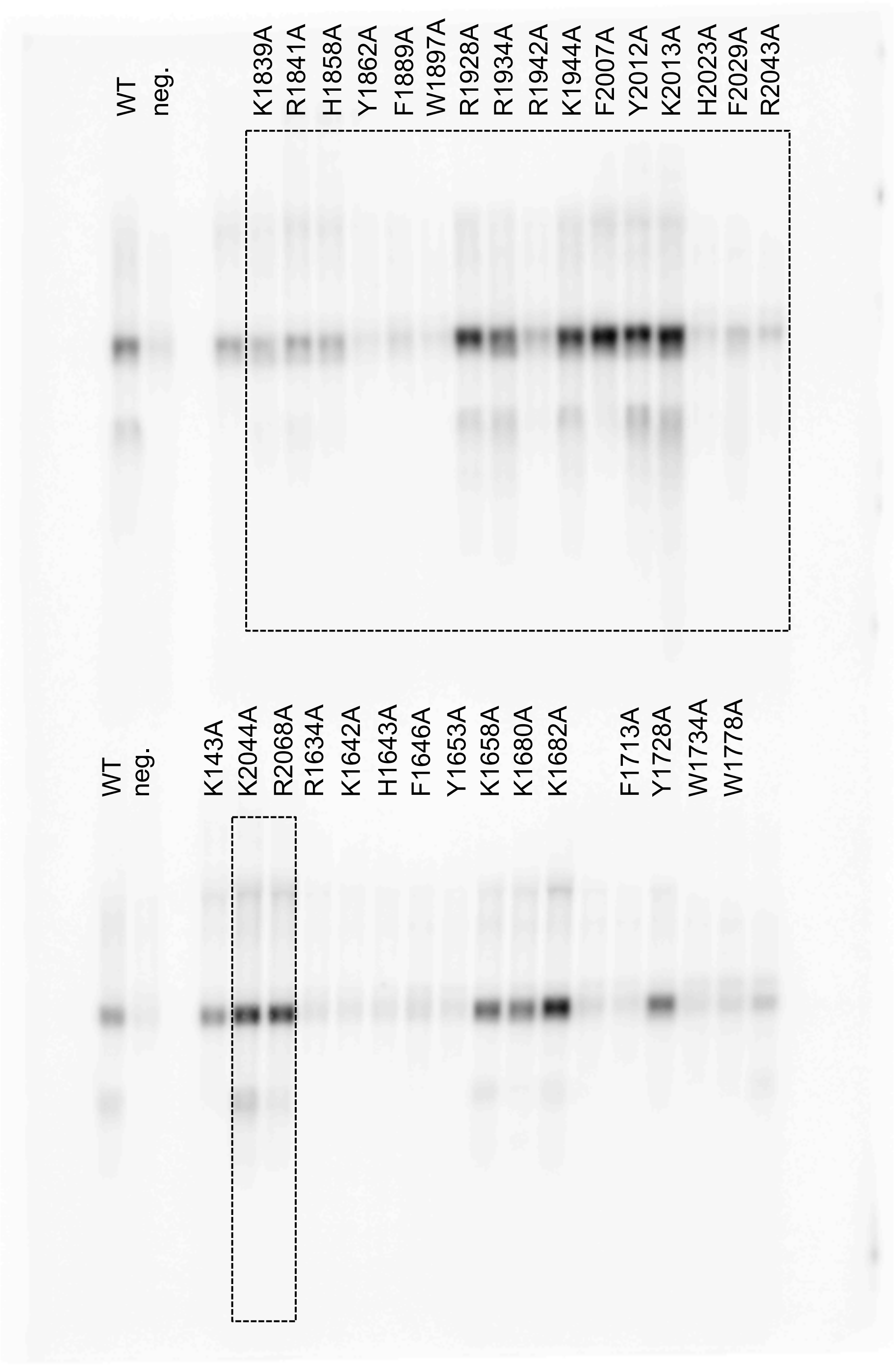

### Methylene blue staining of membrane 2

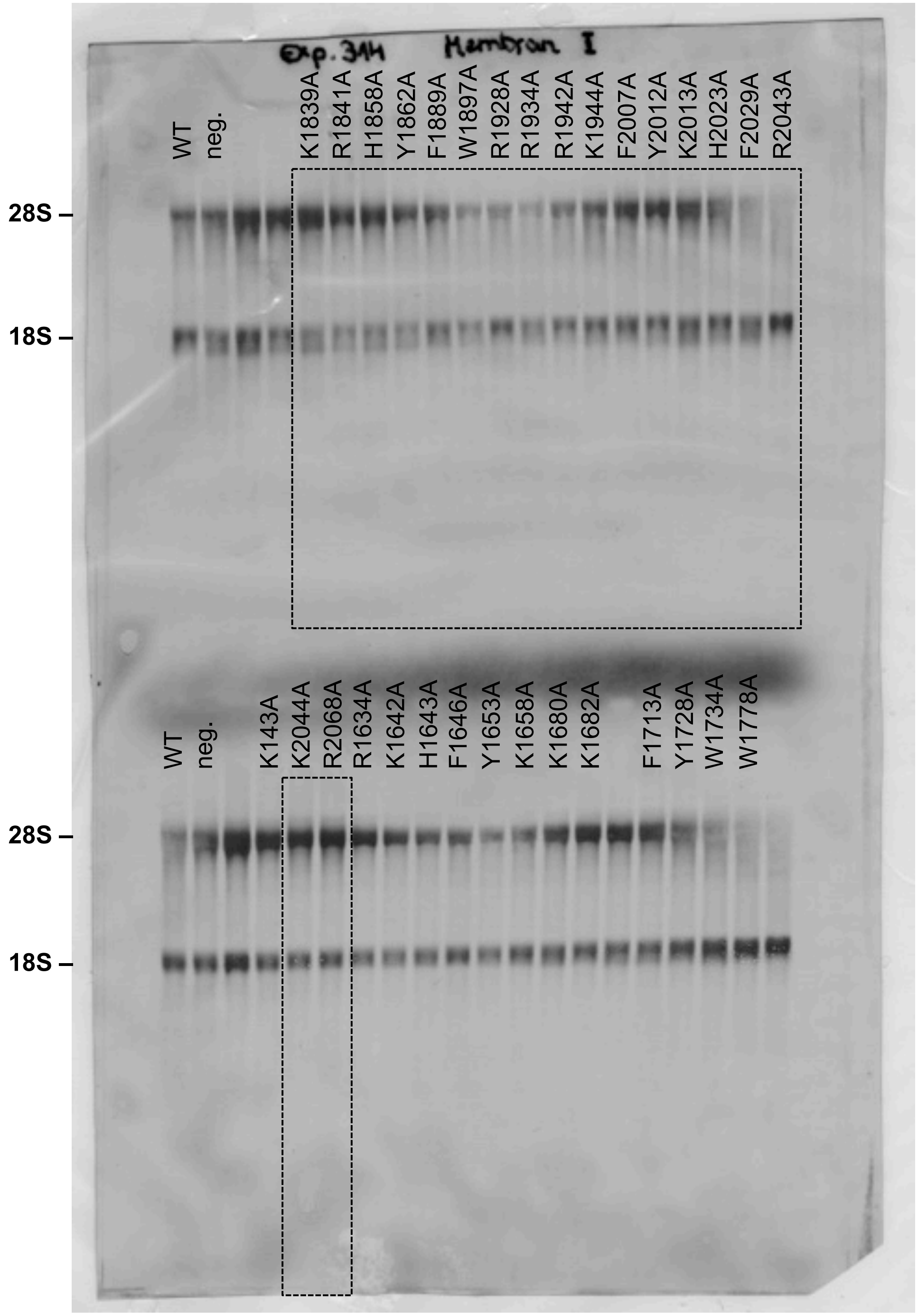

### Northern Blot 3

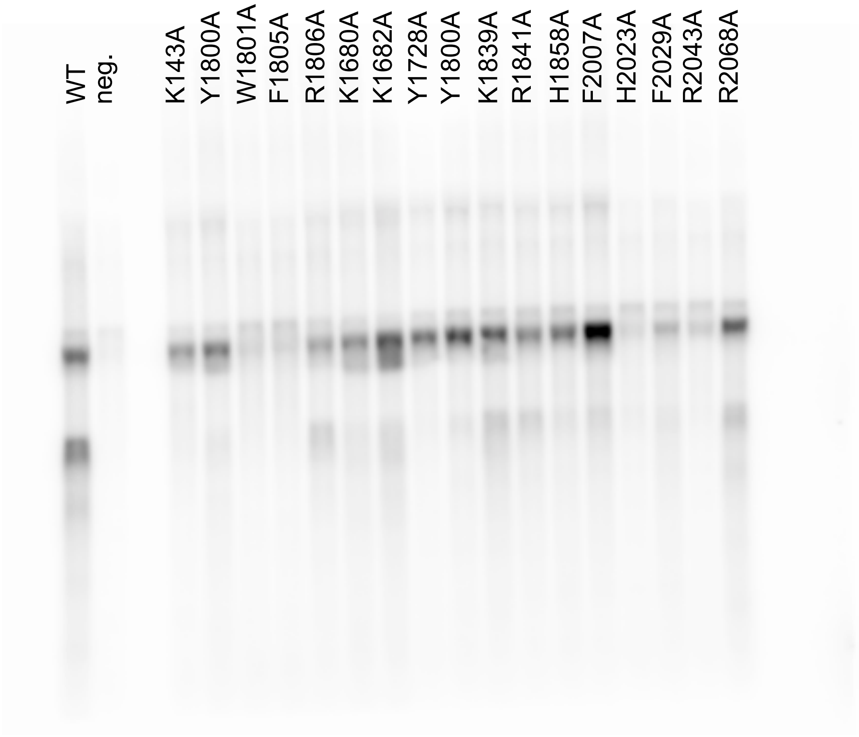

### Methylene blue staining of membrane 3

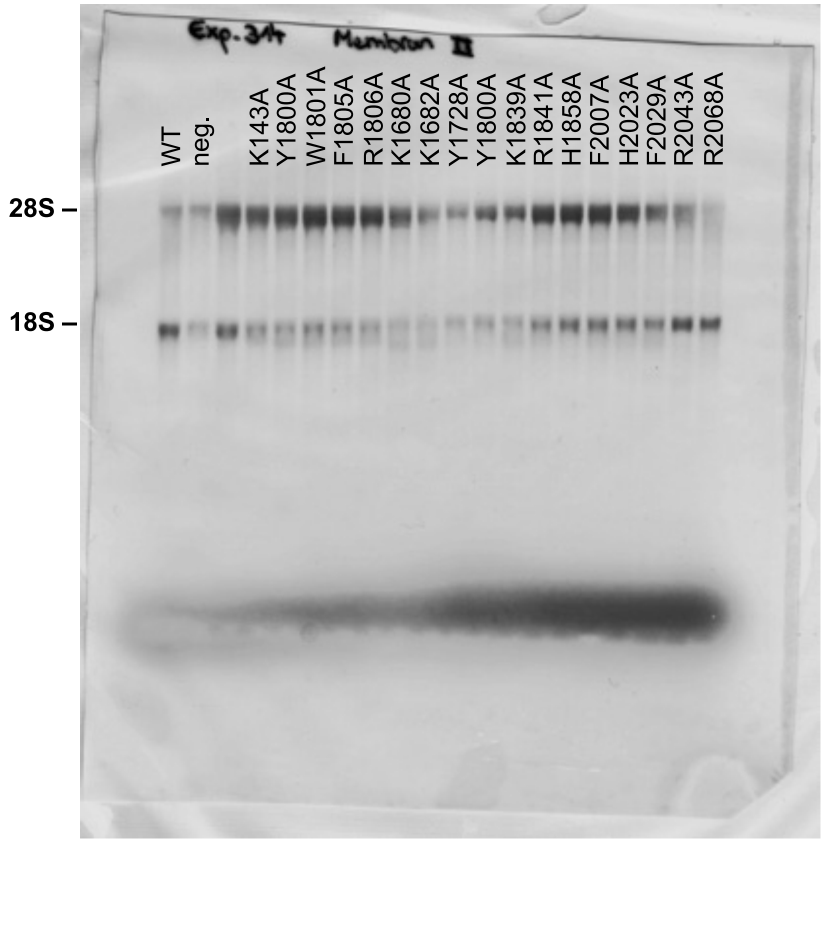

### Northern Blot 4

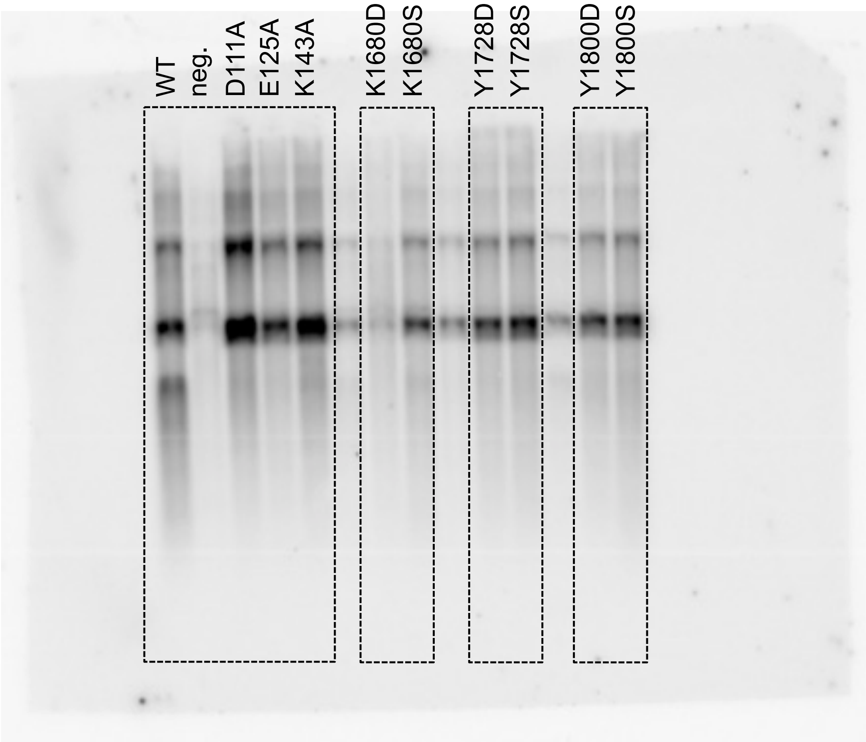

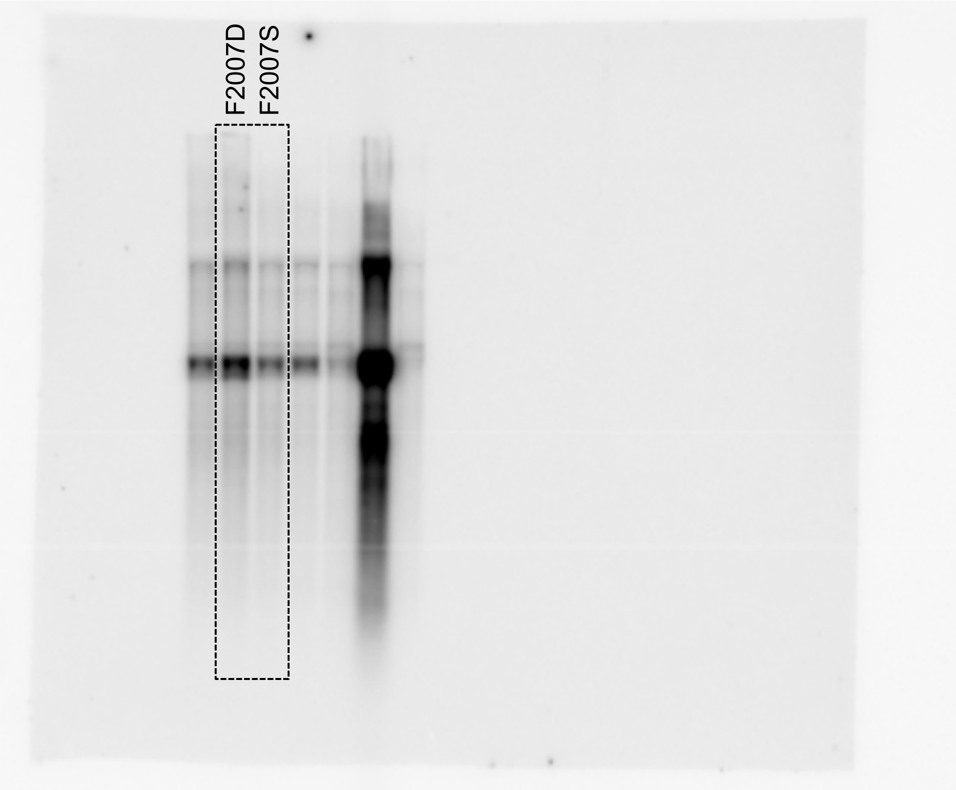

### Methylene blue staining of membrane 4

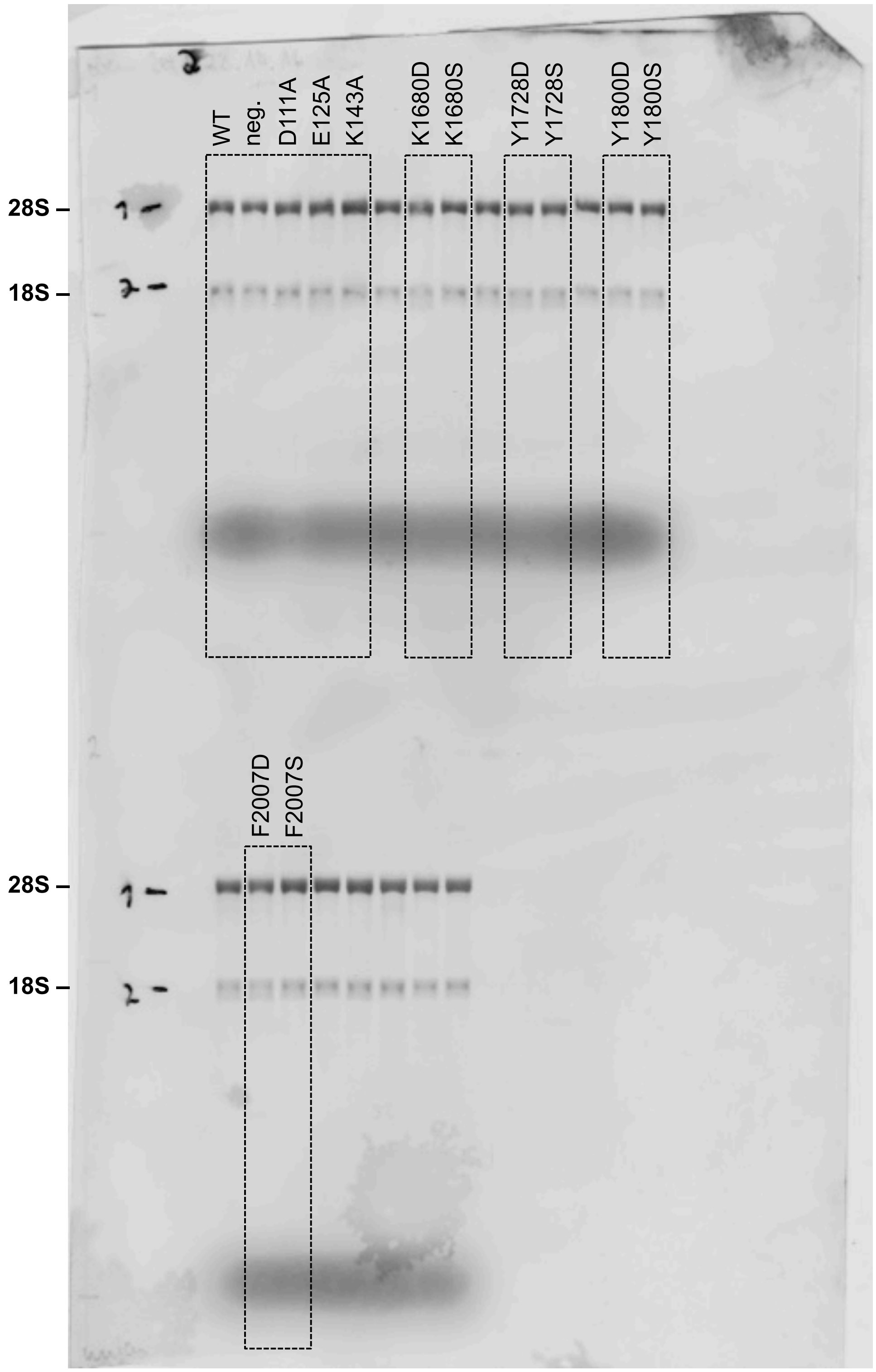

### Northern Blot 5

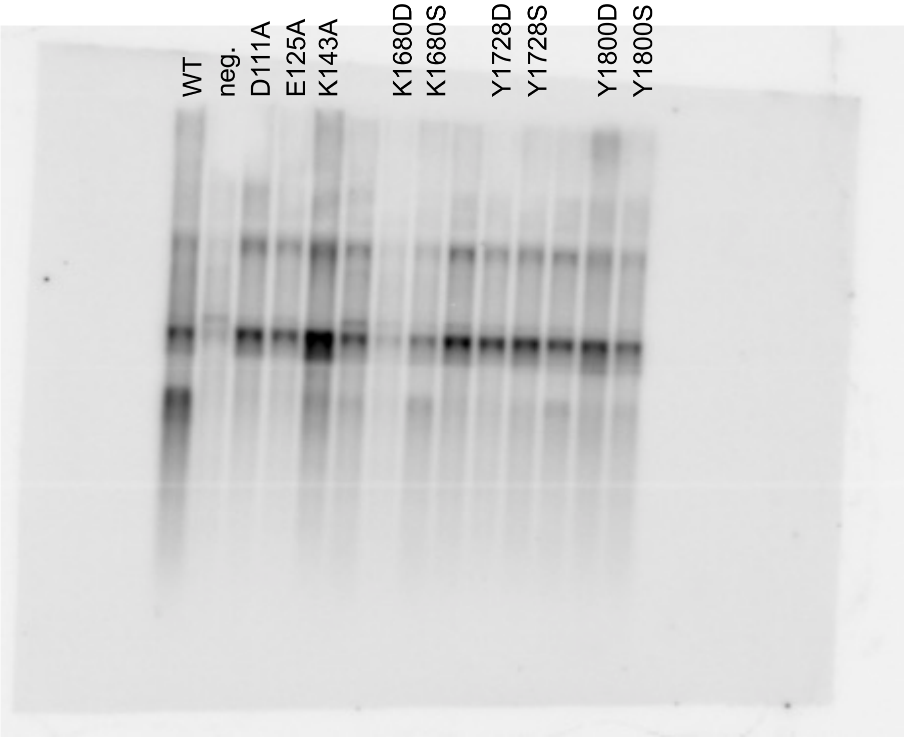

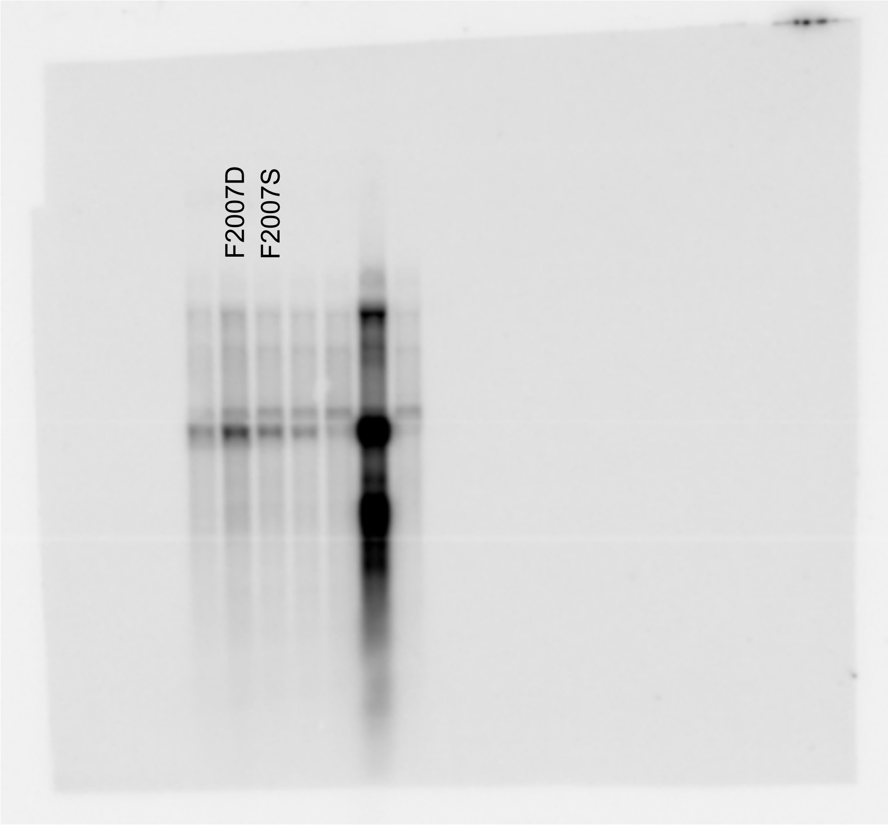

### Methylene blue staining of membrane 5

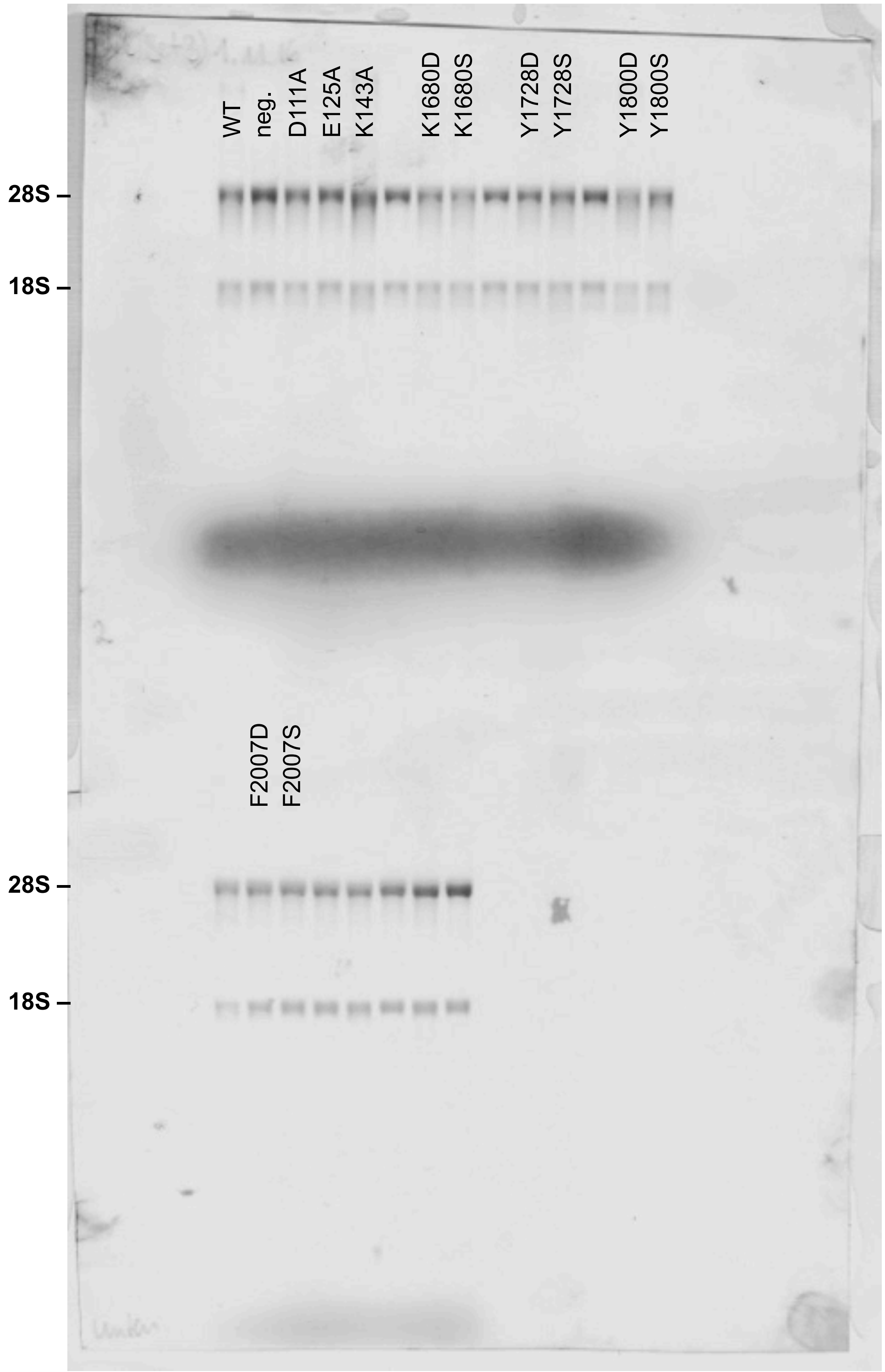

### Western Blots

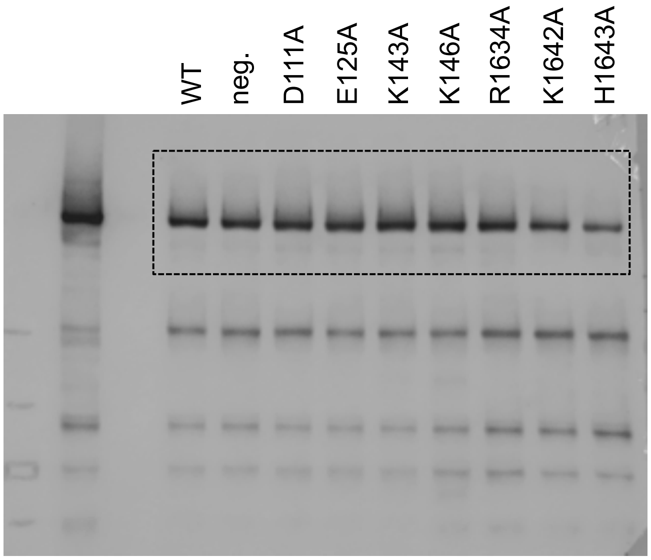

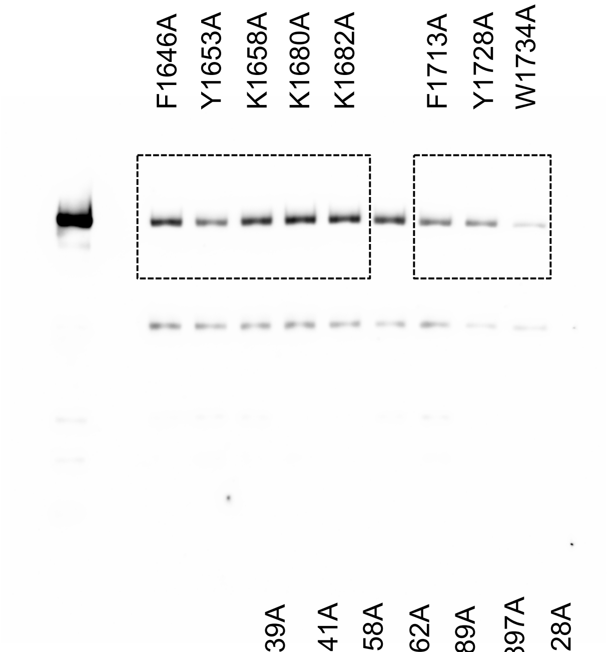

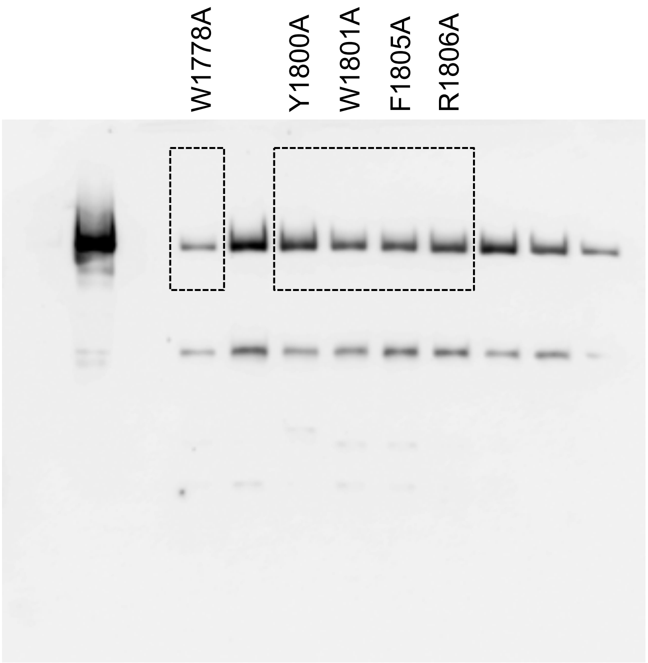

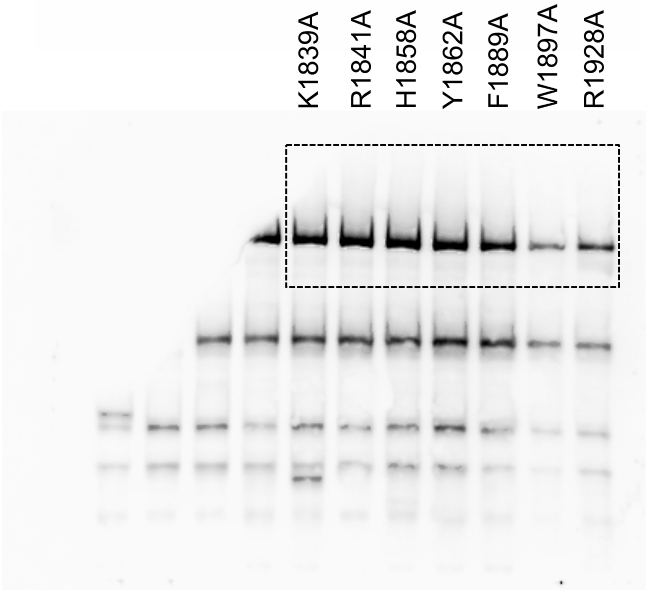

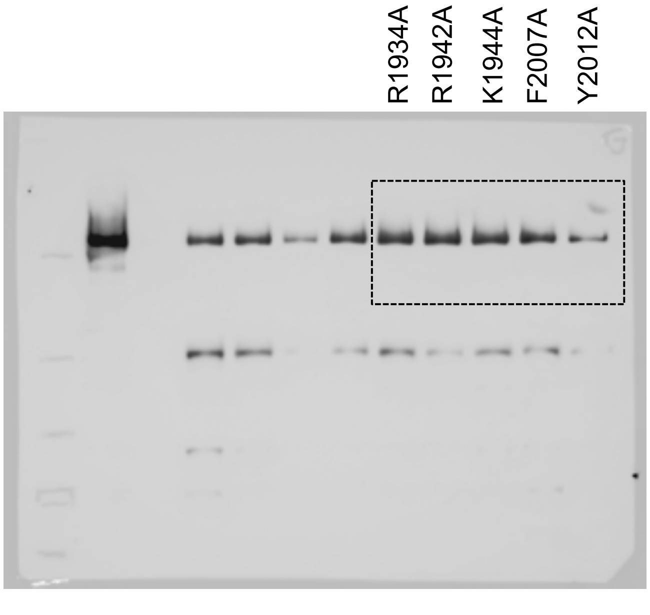

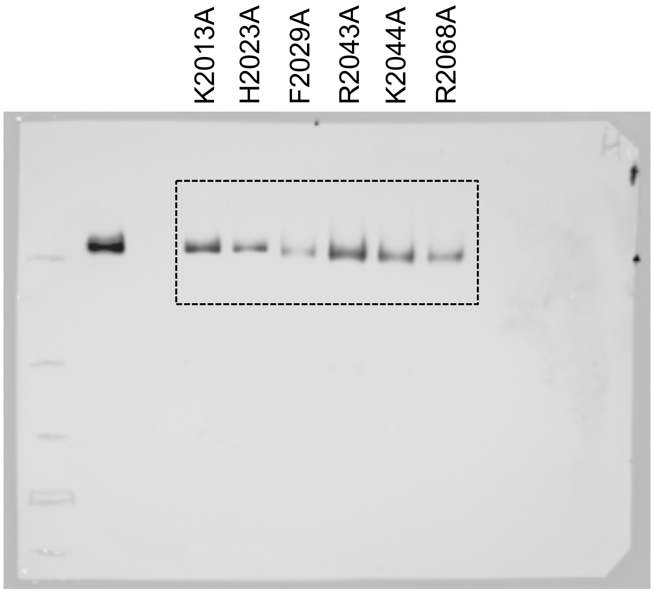
